# Bridging Gene Expression and Morphology: A Cell Size Score and Its Applications Across Multiple Diseases and Physiological Contexts

**DOI:** 10.64898/2026.06.28.733694

**Authors:** Xiangwen Ji, Qinghua Cui

**Affiliations:** Department of Cardiology and Institute of Vascular Medicine, State Key Laboratory of Vascular Homeostasis and Remodeling, Peking University Third Hospital, 49 Huayuanbei Road, Beijing 100191, China; School of Sports Medicine, Wuhan Sports University, No. 461 Luoyu Rd., Hongshan District, Wuhan 430079, Hubei Province, China; Department of Biomedical Informatics, State Key Laboratory of Vascular Homeostasis and Remodeling, School of Basic Medical Sciences, Peking University, 38 Xueyuan Rd, Beijing, 100191, China

## Abstract

Cell size is a critical morphological parameter determining cellular functional homeostasis, yet existing large-scale transcriptomic databases lack direct cell size measurement data. By integrating high-resolution immunofluorescence images with transcriptomics, we identified 457 genes significantly correlated with cell area. Based on these findings, we developed an algorithm, Cell Size Score (CSS), to predict cell size from gene expression profiles. Validation across multiple independent datasets, including human cell lines, mouse models, and single-cell spatial transcriptomics, confirmed that CSS accurately predicts cell size. Furthermore, we observed a significant positive correlation between CSS and broad-spectrum chemotherapy drug resistance, suggesting that increased cell volume confers survival advantages to cancer cells. Moreover, CSS analysis of aging revealed sex-dependent, tissue-specific patterns of change, wherein male adipose and cardiac tissues exhibited progressive hypertrophy with age, while female reproductive organs showed significant atrophy. Additionally, CSS significantly increased in skeletal muscle after exercise, indicating that this metric can capture dynamic physiological adaptation processes. This study establishes a bridge between transcriptomics and cell morphology, providing novel insights into retrospectively analyzing the role of cell size in pathological and physiological processes such as cancer and aging using existing omics data, as well as understanding the molecular mechanisms underlying cell size regulation.

## Introduction

Cell size is one of the most fundamental morphological parameters in living systems, and its precise regulation is crucial for maintaining cellular functional homeostasis. From unicellular to multicellular organisms, abnormal changes in cell volume are often closely associated with pathological conditions. For example, in cancer, the loss of cell size uniformity is one of the hallmarks of malignancy^1^. Meanwhile, cellular hypertrophy and giant cell formation are common markers of malignant transformation^2^. During aging, alterations in cell size reflect the decline in metabolic function and tissue remodeling^3^. In developmental biology, cell size determines organ morphology and the efficiency of intercellular communication^4,5^.

Quantification of cell size relies on morphological techniques such as microscopy, fluorescence-activated cell sorting (FACS), or Coulter counters. In recent years, Cell Painting^6,7^ and cell segmentation in high-resolution spatial transcriptomics^8–10^ have provided high-throughput cell size measurements at the single-cell level. However, existing large-scale bulk transcriptome and single-cell transcriptome databases, such as The Cancer Genome Atlas^11^ (TCGA) and the Tabula Sapiens^12^ project, do not contain direct cell size measurement data. In the upstream experiments of single-cell RNA sequencing (scRNA-seq), FACS is widely used for quality control of cell suspensions and enrichment of target cells. By measuring forward scatter (FSC) and side scatter (SSC), FACS can remove cell debris, dead cells, and doublets. However, these upstream physical characteristics are often ignored in subsequent bioinformatics analyses. Downstream data analysis relies primarily on intracellular gene expression profiles for clustering and annotation, rarely incorporating cell size information. This separation of morphological and transcriptomic data limits the possibility of retrospective analysis of cell size as a biomarker in existing omics data.

Cell size is a biological feature actively controlled by complex transcriptional regulatory networks. For example, studies have revealed the master regulatory role of the PI3K/Akt/mTOR pathway in cell volume. mTORC1 phosphorylates and activates S6K1, thereby promoting ribosome biogenesis and enhancing overall translational capacity. Additionally, mTORC1 performs multi-site phosphorylation by recognizing the TOS motif on 4EBP1, forcing it to release the translation initiation factor eIF4E, thereby relieving the restriction on mRNA cap-dependent translation and providing an additive effect for increasing cell volume^13,14^. However, sustained activation of mTORC1 may negatively feedback inhibit insulin receptor substrate (IRS) through the substrate Grb10, thereby weakening PI3K and Akt signaling^15^. The relationship between cell size and the transcriptome is not a simple linear scaling. The expression levels of most genes increase proportionally with cell volume to maintain concentration homeostasis^16^. But cells have evolved a differential scaling mechanism, that is, some genes (such as cell cycle inhibitors) exhibit sub-scaling, with their concentrations diluted as volume increases. Meanwhile, other genes (such as cell cycle activators, proteasomes) exhibit super-scaling, with concentrations rising sharply^17,18^. We hypothesize that the cell transcriptome may contain molecular features sufficient for quantitative inference of cell size, making it possible to reconstruct cell size using existing massive omics data.

In this study (**Figure 1A**), we performed high-throughput transcriptome screening centered on cell size and developed a transcriptome-based algorithm, Cell Size Score (CSS), aimed at quantitatively predicting cell size from gene expression data. After validating the effectiveness of CSS using multiple independent validation sets, we further explored its biological and clinical significance. We found that CSS is significantly positively correlated with resistance to broad-spectrum chemotherapeutic agents, suggesting the translational value of cell size in predicting drug response and patient survival prognosis. In aging research, CSS revealed sex-dependent, tissue-specific patterns of cell size alteration. Finally, we extended the use of CSS in the field of exercise physiology, demonstrating the broad application potential of this metric.

**Figure 1.**
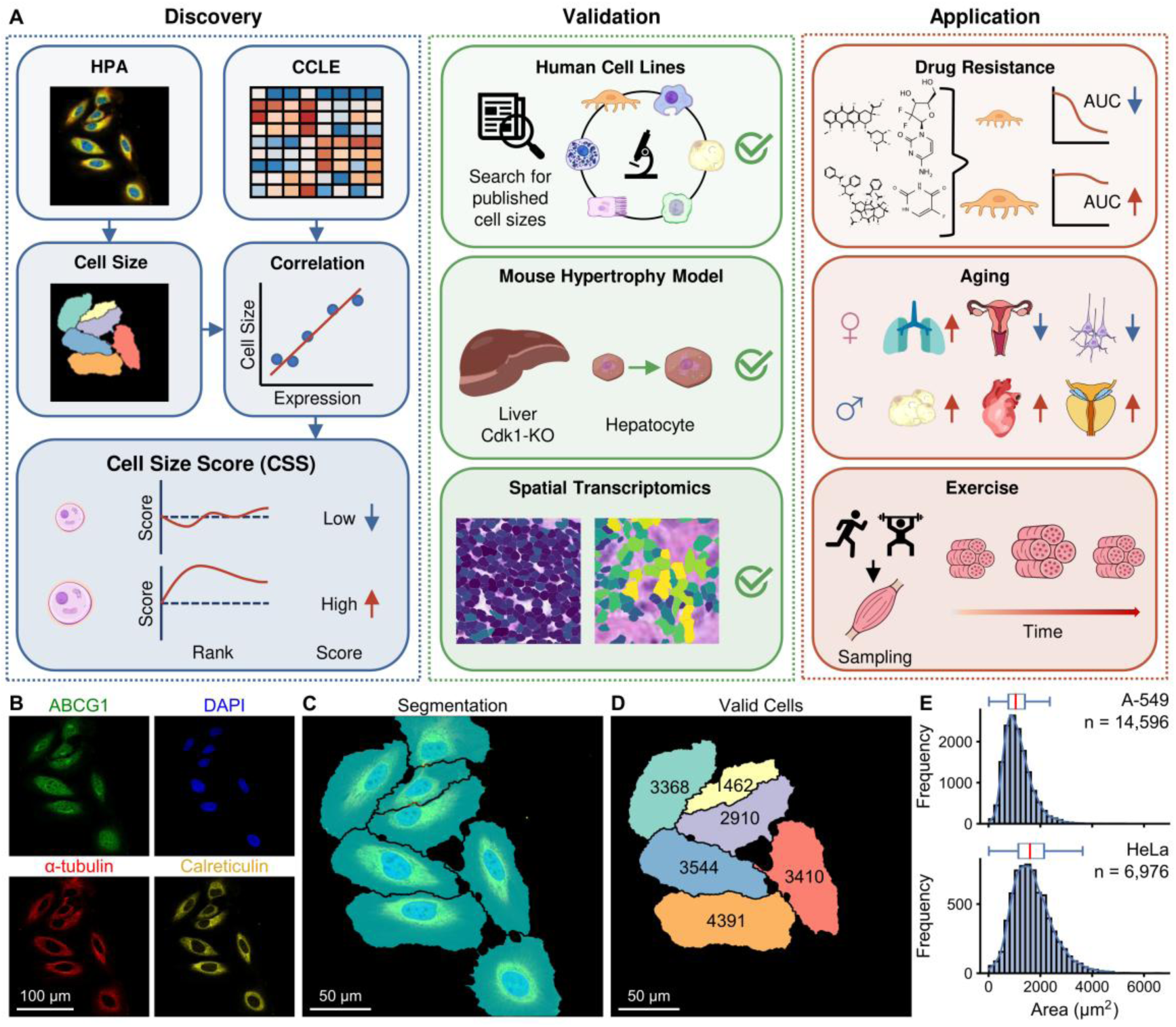
Research pipeline overview and quantification of cell line size. (A) An overview of this research. Original immunofluorescence (IF) images in panel (A-C) were obtained from the HPA (https://images.proteinatlas.org/31470/1046_D11_2_blue_red_green_yellow.jpg). The schematics of several cells and organs are sourced from SciDraw (https://scidraw.io). CCLE: Cancer Cell Line Encyclopedia. (B) Representative multiplexed IF images of cells. The panels display individual channels for ABCG1 (green), nuclei (DAPI, blue), microtubules (α-tubulin, red), and endoplasmic reticulum (Calreticulin, yellow). Scale bar = 100 μm. (C) Cell segmentation of the IF image in (B), overlaid on the composite fluorescence image. Scale bar = 50 μm. (D) Identification of valid cells. Non-border-touching cells are isolated and pseudo-colored. The superimposed numbers indicate the quantified area (μm²) of each cell. Scale bar = 50 μm. (E) Histograms and kernel density estimation (KDE) curves illustrating the distribution of cell area (µm²) for A-549 (top, n = 14,596) and HeLa (bottom, n = 6,976) cell lines. The horizontal box plots above each histogram summarize the descriptive statistics of the population: the center line denotes the median area, the left and right edges of the blue box indicate the 25th and 75th percentiles, respectively, and the whiskers extend to the most extreme values within 1.5 times the interquartile range.

## Results

### Definition and Validation of the Transcriptome-Based Cell Size Score

Accurate quantification of cell size is a prerequisite for our study. To comprehensively and precisely determine the subcellular localization of various human proteins, the Human Protein Atlas (HPA) project employed high-throughput and standardized experimental workflows to systematically capture and analyze ∼100,000 high-resolution immunofluorescence (IF) images (**Figure 1B**), thereby providing a solid data foundation for elucidating the spatial mapping relationships between protein function and cell structure^19,20^. Beyond the original purposes of these images, thanks to this unified experimental workflow, we can utilize these images to obtain stable and comparable cell area information across different cell lines. For each image, we can obtain the pixels occupied by cells through cell segmentation (**Figure 1C**) and convert them to area using scale bars (**Figure 1D**). By integrating results from 63,475 images and 519,632 cells, the median cell areas of 22 cell lines (Supplementary Table S1) were calculated, constituting our discovery dataset. Cell areas vary significantly among different cell lines (**Figure 1E**, Supplementary Figure S1). REH and Jurkat exhibit highly compact area distributions with lower median values. In contrast, HeLa and GAMG show larger median areas and wider interquartile ranges, reflecting greater variability in cell morphology within these populations.

The Cancer Cell Line Encyclopedia (CCLE) comprehensively records transcriptomic information of thousands of human tumor cell lines, providing important data support for cancer basic research and the development of anti-tumor drugs^21^. Based on the above data, using cell lines as a bridge, we were able to study the transcriptomic features of cell size. We evaluated the correlation between the whole transcriptome and cell area, identifying 457 significantly (false discovery rate, FDR < 0.05) positively correlated and 120 significantly negatively correlated genes (**Figure 2A**, Supplementary Table S2). Interestingly, among the most significantly positively correlated genes, we identified candidate genes with potential cell size associations such as GPX8, LRRC49, and SNX7. Recent studies indicate that GPX8 can inhibit apoptosis and promote survival and invasive growth of tumor cells^22^. LRRC49 has been confirmed to participate in microtubule polymerization, which is a key physical basis necessary for cytoskeletal expansion^23^. Moreover, SNX7 inhibits both cellular autophagy and apoptosis^24,25^. Taking GPX8 as an example, its expression level shows a strong positive correlation with cell area (**Figure 2B**, Spearman’s correlation, ρ = 0.92, FDR = 5.3×10^-5^). These findings suggest that our screening effectively captured genes with biological significance related to cell size.

**Figure 2.**
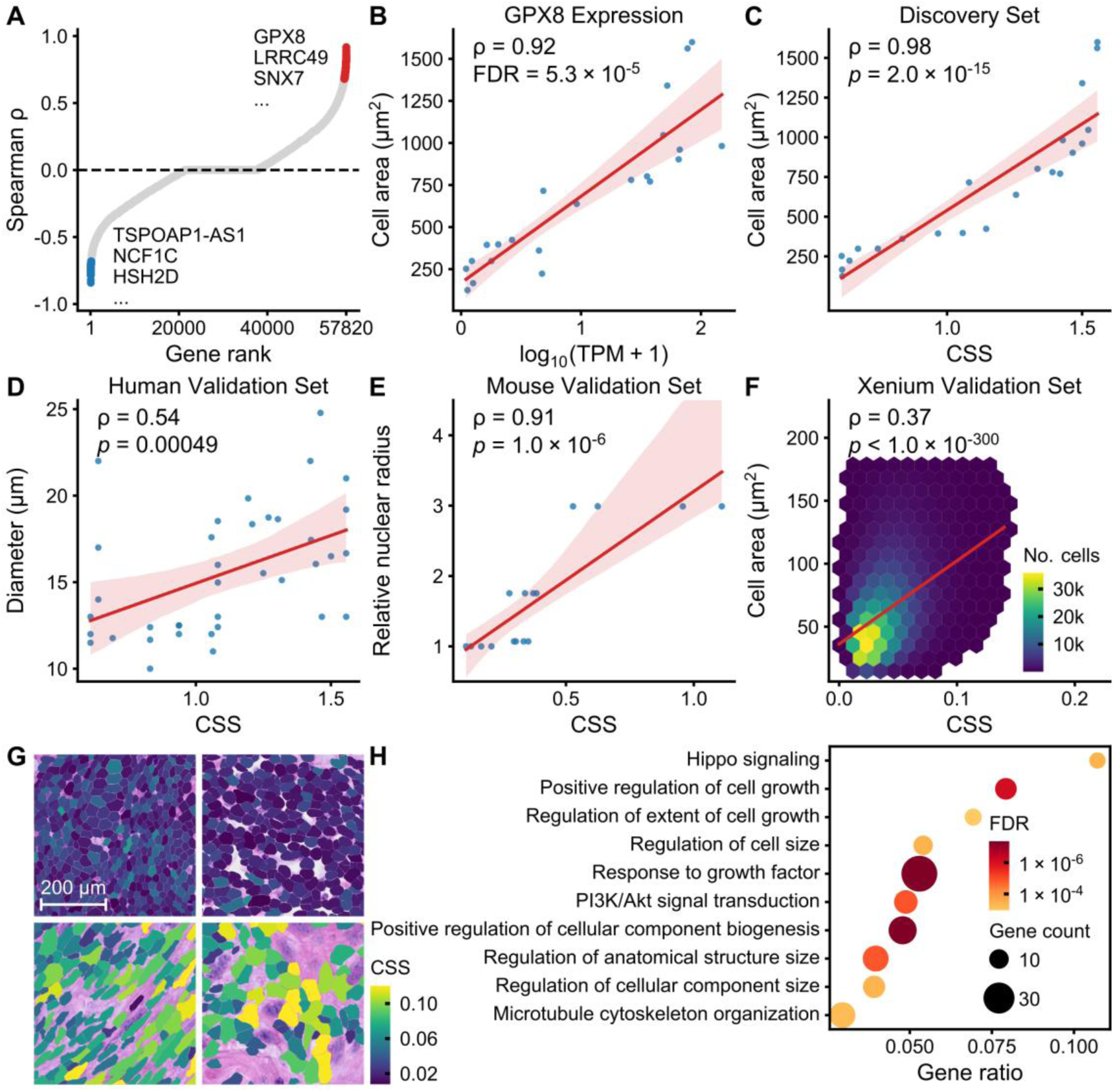
Construction and validation of the Cell Size Score (CSS). (A) Spearman correlation (ρ) ranking of 57,820 genes against cell area. Red and blue dots indicate the significant (false discovery rate, FDR < 0.05) positively and negatively correlated genes, respectively. (B) The correlation between GPX8 expression and cell area (μm^2^). The red line represents the linear regression fit, and the shaded area denotes the 95% confidence interval (same for C-E). TPM: transcripts per million. (C) The correlation between the derived CSS score and cell area in the discovery cohort. (D-E) Validation of the CSS in independent datasets, showing significant correlation with cell diameter (μm) in a human cohort (D) and relative nuclear radius in a mouse cohort (E). (F) Correlation between CSS and segmented cell area (μm^2^) across 732,704 cells using the Xenium spatial transcriptomics platform. Color intensity reflects cell count. (G) Spatial visualization of representative tissue sections. Segmented cell boundaries are filled with colors representing the CSS. Cells with a smaller area tend to have a smaller CSS and vice versa. All the fields of view have the same scale bar = 200 μm. (H) Bubble plot of Gene Ontology (GO) enrichment analysis for genes constituting the CSS. Bubble size indicates gene count, and color gradient represents the FDR.

Based on these results, we constructed CSS to quantitatively assess relative differences in cell size (Methods). In the discovery dataset, CSS shows a high positive correlation with cell area (**Figure 2C**, ρ = 0.98, *p* = 2.0×10^-15^). To test the accuracy of this score, we performed validation in independent datasets. First, we manually searched and curated 37 reports of diameters of human cell lines from literature and databases^26–30^, involving 21 cell lines (Supplementary Table S3). We calculated the correlation between cell diameter and CSS of the corresponding cell lines calculated based on CCLE. Results showed that CSS exhibited strong correlation with cell diameter in this validation set (**Figure 2D**, ρ = 0.54, *p* = 4.9×10^-4^). Unlike the highly comparable cell sizes obtained from the unified experimental workflow of HPA in the discovery dataset, cells from literature were under different culture and measurement conditions, resulting in high heterogeneity of results. For example, five studies^27–31^ reported different cell diameters for MCF-7, ranging from 12.41 to 18.53 μm. Therefore, we believe that achieving ρ=0.54 for CSS is already quite good. Miettinen et al.^32^ specifically knocked out the Cdk1 gene in mice to make hepatocytes grow continuously without division, thereby constructing four types of cells with different nuclear radii (positively correlated with cell size) and performing RNA sequencing. In this mouse validation set, CSS was highly correlated with relative nuclear radius (**Figure 2E**, ρ = 0.91, *p* = 1.0×10^-6^). This cross-species consistency reflects the accuracy of CSS and suggests the evolutionary conservation of transcriptomic programs determining cell size.

However, the above datasets are all bulk transcriptomes, reflecting average cell sizes in tissues. To further validate CSS at single-cell resolution, we applied it to a cervical cancer Xenium spatial transcriptomics dataset, which contained 732,704 cells after filtering. Subcellular-resolution Xenium provides cell segmentation based on cell membrane (using ATP1A1, CD45, and E-Cadherin) or cytoplasmic (using 18S ribosomal RNA) markers, naturally carrying cell size information. Due to systematic morphological errors introduced by physical deformation during tissue section preparation, mutual compression between cells in dense microenvironments, and area shrinkage caused by non-equatorial sectioning, cell area often cannot perfectly equal true cell size. Additionally, the Xenium technology measures a panel of ∼5,000 genes, resulting in only 133 genes (29.1% of complete genes) available for calculating CSS, and the results are also affected by data sparsity. Despite these challenges, CSS scores at the single-cell level still showed ideal positive correlation with segmented cell areas (**Figure 2F, G**, ρ = 0.37, *p* < 1.0×10^-300^).

If CSS truly represents cell size, its constituent genes should be highly correlated with known growth regulatory pathways. As expected, Gene Ontology (GO) enrichment analysis showed (**Figure 2H**) that the genes we screened were significantly enriched in terms related to cell size and cell growth, including positive regulation of cell growth, regulation of extent of cell growth, regulation of cell size, response to growth factor, regulation of anatomical structure size, etc. The results also indicated the important roles of the Hippo signaling pathway and the PI3K/Akt signaling pathway. The Hippo pathway is a classic mechanical stress transduction and organ/cell size control network^33,34^. Meanwhile, the PI3K/Akt cascade is the metabolic hub driving biomass accumulation and protein synthesis, providing substrate support for physical enlargement of cells^13,35–37^. It should be noted that directly using these enriched GO terms to predict cell size does not yield stable results superior to CSS (Supplementary Table S4), suggesting that existing knowledge can only explain part of the molecular mechanisms of cell size regulation.

### Cell Size Shows Significant Positive Correlation with Broad-Spectrum Chemotherapy Resistance

As a basic cellular morphological feature, the systematic impact of cell size on pharmacological responses remains insufficiently clarified. For individual organisms, larger body size confers greater energy reserves and stronger environmental adaptability^38,39^. We hypothesize that this principle also applies to cancer cells, meaning larger cancer cells are more likely to survive under chemotherapeutic pressure. The Cancer Therapeutics Response Portal (CTRP) provides large-scale data on drug sensitivity of cancer cell lines to hundreds of small molecule compounds, aimed at helping researchers discover potential cancer treatment targets and biomarkers^40^. To investigate the global relationship between cell size and drug sensitivity, we performed correlation analysis between CSS and CTRP drug sensitivity data. Strikingly, in the vast majority of drugs tested, the area under the dose-response curve (AUC), representing cellular drug resistance, showed broad and significant (Spearman test, FDR < 0.05) positive correlations with CSS (**Figure 3A**, Supplementary Table S5). A total of 407 drugs showed a significant positive correlation (with the maximum correlation coefficient being ρ = 0.64), while only 26 drugs showed a significant negative correlation (with the minimum ρ = -0.30). This positive correlation across different drug classes strongly suggests that increased cell volume may endow cells with non-specific, broad-spectrum endogenous drug resistance mechanisms. For instance, we analyzed five first-line clinical chemotherapeutic agents (doxorubicin, oxaliplatin, gemcitabine, paclitaxel, and fluorouracil) with different pharmacological mechanisms. Results (**Figure 3B-F**) were highly consistent with global trends: among these five anti-tumor drugs, cell size maintained strong positive correlations with AUC (Spearman ρ ranging from 0.42 to 0.48, all *p* < 10^-33^).

**Figure 3.**
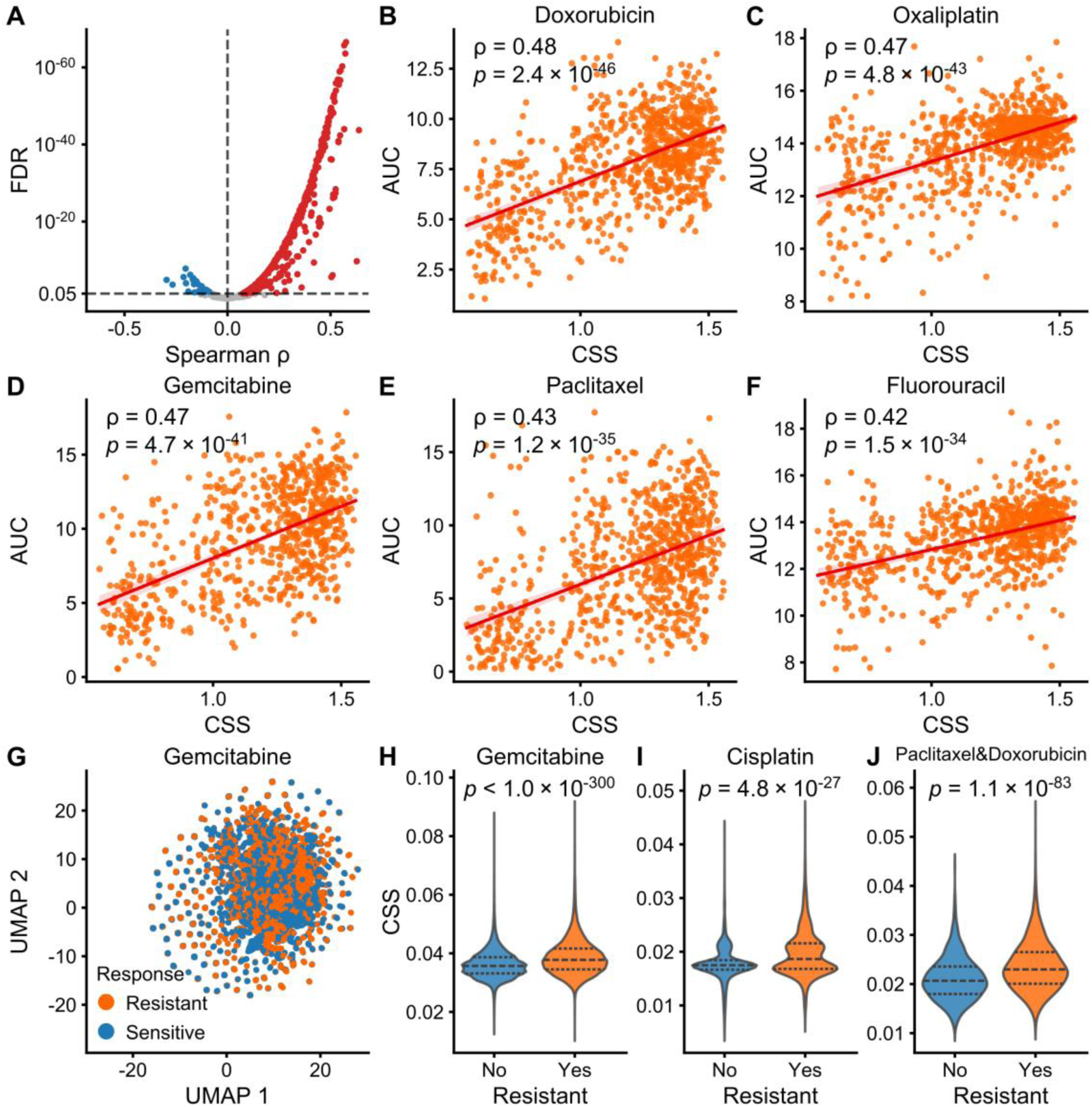
Cell Size Score (CSS) serves as a robust predictor of broad-spectrum drug resistance. (A) The Spearman correlation between the CSS and the area under the dose-response curve (AUC) across all screened compounds. The x-axis represents the Spearman correlation coefficient (ρ), and the y-axis represents the log-transformed false discovery rate (FDR). Red and blue dots indicate compounds with a significant (FDR < 0.05) positive and negative correlation, respectively. (B-F) Correlations between CSS and AUC for five representative first-line chemotherapeutics: (B) Doxorubicin, (C) Oxaliplatin, (D) Gemcitabine, (E) Paclitaxel, and (F) Fluorouracil. The red line represents the linear regression fit. (G) Single cell UMAP visualization of cell clustering based on Gemcitabine drug response. (H-J) CSS distribution between resistant and sensitive cells for (H) Gemcitabine, (I) Cisplatin, and (J) Paclitaxel & Doxorubicin. The dashed lines within the violins represent the median and the 25th and 75th percentiles. Two-sided Wilcoxon’s rank sum tests are used.

To further validate the relationship between cell size and drug resistance at the single-cell level, we analyzed data from scDrugAtlas, which contains single-cell transcriptomic profiles paired with drug response phenotypes (**Figure 3G**, Supplementary Figure S2). We calculated the CSS for each cell and compared the distribution between drug-resistant and drug-sensitive populations. Notably, Uniform Manifold Approximation and Projection (UMAP)^41^ sometimes fails to cluster cells by drug response. For instance, for Gemcitabine, the UMAP completely failed to distinguish between resistant and sensitive cells (**Figure 3G**). However, the CSS in resistant populations was significantly higher than in sensitive populations across various drugs (Wilcoxon rank-sum test, Gemcitabine, *p* < 1.0×10^-300^; Cisplatin, *p* = 4.8×10^-27^; Paclitaxel & Doxorubicin, *p* = 1.1×10^-83^). This contrast suggests that while traditional clustering methods may fail to identify drug-resistant tumor cell subpopulations, CSS provides a robust metric to capture the morphological basis of drug resistance that is otherwise obscured.

Additionally, Cox proportional hazards regression models showed that among 33 cancer types in The Cancer Genome Atlas (TCGA), increased cell size was associated with adverse overall survival in 11 cancer types, including lower grade glioma (LGG), lung adenocarcinoma (LUAD), pancreatic adenocarcinoma (PAAD), etc. (Supplementary Table S6). No significant opposite results were found. These findings not only validate the hypothesis that larger cells have stronger drug resistance, but also indicate that cell size is expected to serve as a biomarker for assessing and predicting tumor cell sensitivity to broad-spectrum chemotherapy.

### Tissue-Specific Changes in Cell Size During Aging

To investigate the dynamic changes in cell size across different anatomical sites during aging, we used the Genotype-Tissue Expression (GTEx) Project^42^ to assess CSS of various tissues/organs and calculated their correlations with age. We observed that cell morphological changes associated with physiological age exhibit high tissue specificity. In some tissues/organs, including adipose tissue (ρ = 0.12, *p* = 2.0×10^-5^), colon (ρ = 0.20, *p* = 8.0×10^-9^), heart (ρ = 0.07, *p* = 0.030), lung (ρ = 0.11, *p* = 0.0072), and prostate (ρ = 0.15, *p* = 0.021), CSS showed significant positive correlations with age (**Figure 4A-D, F**). This progressive enlargement of cells with increasing age is highly consistent with classic morphological features of cellular senescence, where metabolically active but non-proliferating senescent cells typically undergo pathological hypertrophy^43^. Conversely, CSS in neural (tibial nerve) tissues showed a slight but significant negative correlation with age (**Figure 4E**, ρ = -0.13, *p* = 0.0012), which may reflect age-related neural atrophy and degeneration.

**Figure 4.**
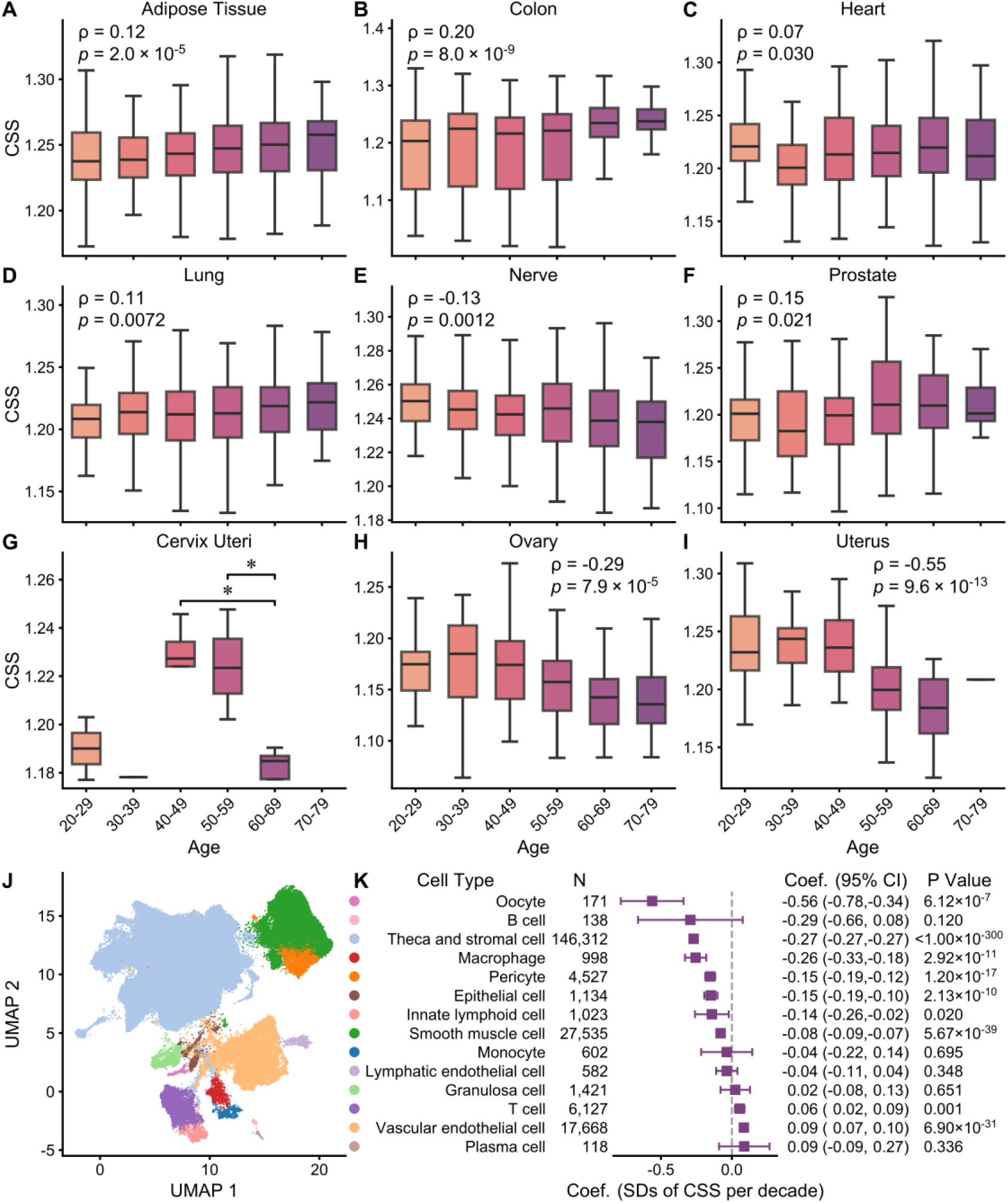
Age-dependent alterations of Cell Size Score (CSS) across multiple human tissues. (A-I) The distribution of CSS across different age decades (20-29 to 70-79 years) in (A) adipose tissue, (B) colon, (C) heart, (D) lung, (E) nerve, (F) prostate, (G) cervix uteri, (H) ovary, and (I) uterus. The center line of each box represents the median, box limits represent the 25th and 75th percentiles, respectively, and the whiskers extend to the most extreme values within 1.5 times the interquartile range. Spearman correlation coefficients (ρ) and p values are annotated for (A-F, H, and I). For the cervix uteri (G), two-sided Wilcoxon’s rank sum tests are used, \**p* < 0.05. (J) UMAP visualization of distinct cell clusters in human ovaries. The correspondence between color and annotated cell type is shown in (K). (K) The correlation between CSS and age for each ovarian cell type. Coef. denotes the coefficient for age in the linear model (change in the standard deviation [SD] of CSS per decade), with data batches included as covariates. N: number of cells; CI: confidence interval.

Most significantly, female reproductive organs exhibited strong reduction in cell size with increasing age. Ovary (**Figure 4H**, ρ = -0.29, *p* = 7.9×10^-5^) and uterus (**Figure 4I**, ρ = -0.55, *p* = 9.6×10^-13^) both showed strong negative correlations, with absolute values of correlation coefficients higher than other tissues/organs. Additionally, due to small sample sizes for cervix uteri (2 cases in 20-29 group, 1 case in 30-39 group), overall correlation could not be observed. However, statistical testing between groups revealed a statistically significant decline in CSS after menopause (60-69 years) (**Figure 4G**, Wilcoxon rank-sum test, *p* < 0.05, compared to both the 40-49 and 50-59 groups). This highlights the tight coupling between endocrine microenvironments and cell morphology.

To further dissect the cellular heterogeneity of this aging process within the ovary, we integrated multiple scRNA-seq datasets of human ovaries across different ages. We identified 14 major cell types (Methods, **Figure 4J**, Supplementary Figure S3A-D) and calculated the CSS for each cell type and evaluated its correlation with age (**Figure 4K**). Strikingly, oocytes, the largest cells in humans, exhibited the strongest negative correlation with age, indicating profound age-related atrophy. Additionally, considering that theca and stromal cells constitute the most abundant cell population in the ovary^44^, their significant negative correlation with age suggests that they are the primary contributors to the overall shrinkage of the aging ovary. In contrast, vascular endothelial cells and T cells showed significant positive correlations with age. This may be because their morphology is relatively less sensitive to the endocrine decline, allowing them to exhibit the typical age-related enlargement characteristic of senescent cells. Interestingly, we observed a significant negative correlation in macrophages (**Figure 4K**). A large proportion of macrophages in young tissues are tissue-resident macrophages, which are characterized by their long lifespan and robust self-renewal capacity. However, in aged tissues, these resident macrophages gradually decline, necessitating the infiltration of blood-derived monocytes into the tissue for replenishment^45,46^. Similarly, we found that the proportion of macrophages significantly decreased in the older age group (>45 years) compared to the younger group (≤45 years) (Supplementary Figure S3E, 52.9% vs. 68.3%). It is well known that macrophages are larger than monocytes, which can also be observed by CSS (Supplementary Figure S3F). These newly converted macrophages likely possess an intermediate size between monocytes and mature macrophages, thereby lowering the average CSS in the older group. Collectively, CSS successfully deciphers the heterogeneous cellular aging trajectories including endocrine-driven atrophy, senescent cell hypertrophy, and immune remodeling within the same organ.

Considering the complex interactions between the endocrine system and aging, we performed further stratified analyses by biological sex for the above non-sex-specific tissues/organs (adipose tissue, colon, heart, lung, and nerve) (Supplementary Figure S4). Interestingly, progressive cell enlargement in adipose tissue (ρ = 0.15, *p* = 1.7×10^-5^) and heart (ρ = 0.11, *p* = 0.0092) was statistically significant only in males (Supplementary Figure S4A, C, F, H), which aligns with epidemiological observations that males are more prone to fat accumulation^47^ and myocardial hypertrophy^48^. Conversely, females exhibited more significant aging-related morphological changes in lung (ρ = 0.18, *p* = 0.018) and neural tissues (ρ = -0.24, *p* = 8.0×10^-4^) (Supplementary Figure S4D, E, I, J). The increase in lung CSS may be related to accelerated loss of collagen and elastin in the lungs of postmenopausal females^49^. Findings in neural tissues also align with the phenomenon that females are more susceptible to tarsal tunnel syndrome^50^. These findings indicate that cellular manifestations of aging, whether hypertrophy or atrophy, are heavily modulated by sex-specific physiological backgrounds.

### Physical Exercise Significantly Increases Skeletal Muscle CSS

Beyond aging, we wondered whether CSS could also indicate cell size alterations in other physiological states. Obviously, physical exercise is a good example. Therefore, we collected 10 independent cohorts, which measured skeletal muscle transcriptome data before and after exercise. Based on these data, we quantified the CSS of skeletal muscle. Across cohorts with different exercise protocols, we consistently observed that CSS of skeletal muscle post exercise was significantly upregulated compared to pre-exercise baseline levels (**Figure 5**). These cross-cohort results further validate the efficacy of the CSS we constructed.

**Figure 5.**
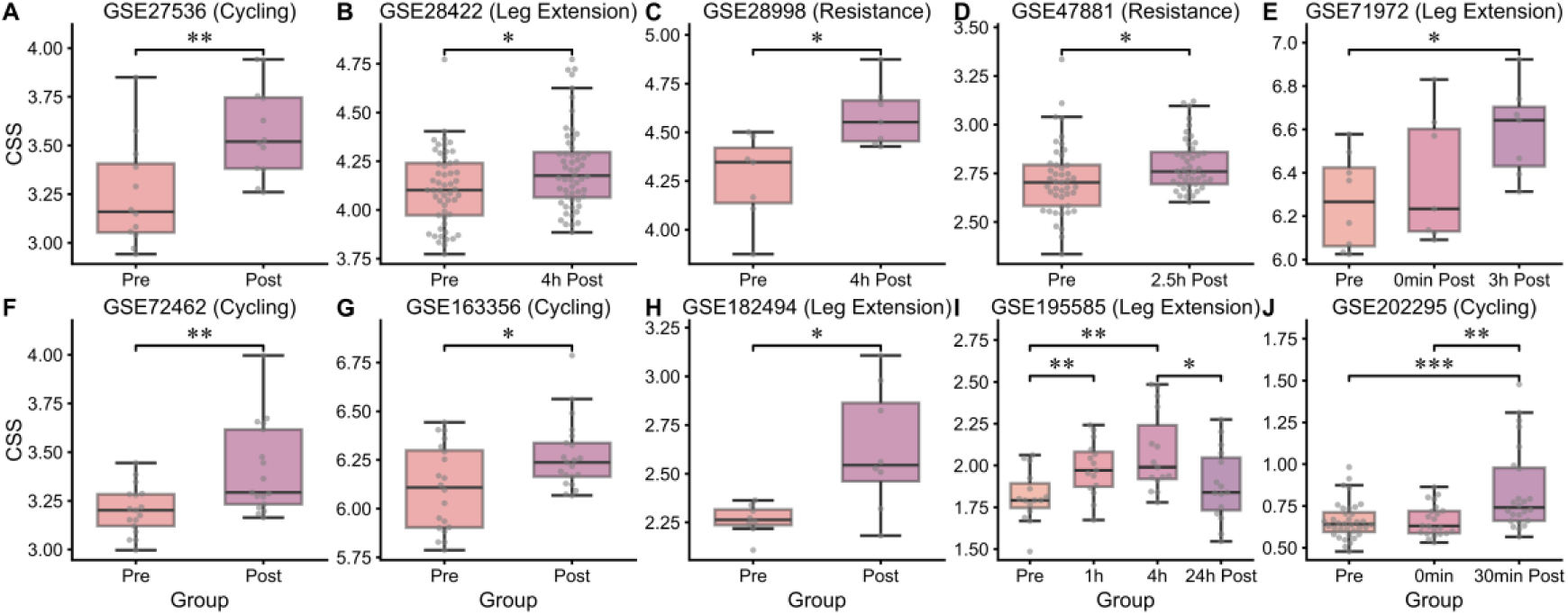
Physical exercise induces a significant increase in the skeletal muscle Cell Size Score (CSS). (A-J) The changes in CSS of skeletal muscle samples before and after physical exercise (Pre vs. Post, including various recovery time points) across 10 independent GEO datasets (GSE27536, GSE28422, GSE28998, GSE47881, GSE71972, GSE72462, GSE163356, GSE182494, GSE195585, and GSE202295). The central line within each box represents the median, the bounds of the box indicate the 25th and 75th percentiles, and the whiskers extend to the most extreme values within 1.5 times the interquartile range. Two-sided Wilcoxon’s rank sum test, \**p* < 0.05, \*\**p* < 0.01, \*\*\**p* < 0.001.

Interestingly, through analysis of datasets containing continuous time points, we found that exercise-induced CSS elevation exhibits significant temporal dynamics. Skeletal muscle CSS showed a progressive increase at 1 hour and 4 hours post-exercise, but declined after 24 hours (**Figure 5I**), suggesting that exercise-induced cell volume changes may involve acute physiological swelling or early compensatory transcriptional responses. Additionally, in two different cohorts (**Figure 5E, J**), CSS changed immediately post-exercise (0 min post) but not significantly, while significant increases appeared subsequently (at 3 h and 30 min, Wilcoxon rank-sum test, *p* < 0.05 and *p* < 0.001, respectively). These results reveal that the response of skeletal muscle cells to physical exercise is a time-sensitive dynamic process, providing new biological insights into understanding exercise-induced skeletal muscle remodeling.

## Discussion

Through high-throughput screening and cross-dataset validation, our study successfully constructed a transcriptome-based CSS and validated its effectiveness across multiple biological scenarios. Gene expression profiles contain abundant morphological information, enabling us to quantitatively reconstruct cell size from large-scale transcriptome databases (such as TCGA and GTEx) in the absence of direct morphology information. This methodological breakthrough provides a new tool for retrospective analysis of the relationship between cell morphology and disease using existing massive omics data, addressing the gap between transcriptomics and morphology.

Enrichment analysis of cell size-correlated genes revealed significant enrichment of the PI3K/Akt/mTOR pathway, which is highly consistent with classical theories on cell growth regulation^36,37^. As the switch of cell volume, mTORC1 drives cell enlargement by promoting ribosome biogenesis and protein synthesis^13,14^. The cross-species conservation of CSS observed in our study (Cdk1 knockout mouse model) reflects the central role of cell volume regulation in evolution. However, our screening also identified a set of genes with significant associations but incompletely characterized functions. Furthermore, although CSS is highly correlated with known pathways, prediction using genes from these pathways directly yields unstable results. These observations suggest that cell size may be an attribute resulting from the cumulative effect of numerous small effects across the genome, with a regulatory network far more complex than currently recognized. Moreover, the causal relationship between transcriptome and morphology may be bidirectional. Changes in cell size can regulate the transcriptome through mechanical stress feedback, with mechanically sensitive transcription factors such as YAP/TAZ serving as typical examples^34^. Therefore, CSS may reflect not only the causes of cell enlargement but also the consequences resulting from it.

This study reveals significant positive correlations between cell size and broad-spectrum chemotherapeutic drug resistance, with potential translational medical significance. This resistance correlation can be explained from a biophysics perspective. According to Fick’s law of diffusion, drug transmembrane flux is proportional to surface area, but cell volume growth far exceeds surface area growth, causing a sharp decline in surface-area-to-volume ratio (S/V Ratio)^51^. This not only limits passive diffusion efficiency of drugs but also dilutes the concentration of drug-target binding by increasing cytoplasmic volume^52^, helping cancer cells survive under chemotherapy pressure. Furthermore, polyploid giant cancer cells (PGCCs) have been discovered, which obtain genomic buffering effects through whole-genome doubling, enhancing resistance to oxidative stress and DNA damage^53,54^. Therefore, CSS may help identify a subpopulation of cancer cells which gain non-specific survival advantages through volume enlargement. This drug resistance, not arising from specific drug target mutations, differs from traditional understanding of resistance mechanisms based on gene mutation typing. We observed that CSS is associated with poor prognosis in multiple cancer types, suggesting that combining genomic features with cell morphological features might enable more precise medical strategies. Existing studies have found that mifepristone can block the embryonic-like developmental programs of PGCCs, preventing them from producing drug-resistant progeny through asymmetric division^55^. Macrolide compounds can block PGCC dormancy signals by inhibiting AXL receptors^56^. These phenomena suggest that sensitization strategies targeting the large cell phenotype may be more effective than targeting specific mutations, such as combination use of mTOR inhibitors or cytoskeleton disruptors.

From the perspective of evolutionary ecology, life history strategies involve a fundamental trade-off between rapid reproduction and robust survival. In natural populations, r-selected species prioritize high reproductive rates and small body size. Conversely, K-selected species invest in large body size and slow reproduction to enhance competitive adaptability and environmental resilience (greater strength, energy reserves, resistance to environmental changes, migratory capacity, etc.)^57^. Notably, this observation is strikingly similar to phenomena in human cellular economy. As confirmed by comprehensive surveys of cell turnover, small, rapidly dividing cells (e.g., erythrocytes and intestinal epithelial cells) dominate in numerical abundance and daily turnover rate (approximately 86%), while large, metabolically specialized cells (e.g., myocytes and adipocytes), despite lower turnover frequency, represent the majority of cellular mass (approximately 75%)^58,59^. This inverse relationship between cell size and abundance suggests that cell populations throughout the human body also follow a size-number trade-off to optimize resource allocation between proliferation and functional capacity. In the context of malignant tumors, we observed that larger tumor cells exhibit enhanced resistance to broad-spectrum chemotherapeutic drugs, indicating that the large cell phenotype confers survival advantages under therapeutic pressure. Interestingly, in survival analysis across 33 cancer types, we consistently found that elevated CSS is associated with poor prognosis in 11 cancers, with no significant evidence of the opposite. This asymmetric result suggests that in the tumor microenvironment, selective pressure favors K-selection strategies. That is, cells with increased volume and enhanced stress resistance achieve superior long-term survival, providing a new perspective for understanding tumor heterogeneity and therapeutic resistance.

Besides cancer, CSS demonstrates great potential in elucidating dynamic changes in normal human cells during physiological and pathological processes. In aging studies, the cell size changes we observed are not uniform but exhibit significant tissue/cell specificity and sex dependence. The severe atrophy of female reproductive organs with age is closely related to tissue degeneration caused by decreased estrogen levels after menopause^49^. In the ovary, our single-cell analysis revealed that this atrophy is predominantly driven by the theca and stromal cells. Notably, vascular endothelial cells and T cells escaped this endocrine-driven atrophy and instead exhibited age-related hypertrophy, indicating that while reproductive organs undergo massive remodeling, some fundamental cellular aging processes still occur in cells that are less dependent on the endocrine system. Intriguingly, we observed a counterintuitive shrinkage in macrophages within the aging ovary. This phenomenon likely reflects a shift in the immune landscape: the gradual loss of long-lived, large tissue-resident macrophages, which are subsequently replaced by smaller, monocyte-derived macrophages recruited from the circulation.

Conversely, the progressive hypertrophy of male adipose and cardiac tissues reflects attempts by these tissues to compensate for decreased function by increasing cell volume during aging, but this compensation ultimately leads to pathological changes. Additionally, the more significant atrophy trends exhibited by tibial neural tissues in females may be related to higher susceptibility of females to tarsal tunnel syndrome^50^. This provides explanations at the cellular morphological level for understanding related diseases, and genes constituting CSS may hide new clues about disease molecular mechanisms.

In the field of exercise physiology, the significant increase in skeletal muscle CSS after exercise further confirms the universality of this indicator. Exercise-induced cell volume enlargement is a complex dynamic process involving osmotic changes and inflammatory responses in the acute phase, as well as increased myofiber protein synthesis in later stages^60,61^. The temporal dynamics we observed, with CSS peaking 1-4 hours post-exercise and returning to baseline within 24 hours, may reflect the temporal lag between transcriptional response and structural remodeling. This suggests that CSS is not only a static morphological indicator but also a biomarker capable of capturing dynamic changes in cellular adaptation processes.

The limitation of this study is that CSS based on bulk transcriptome reflects the average cell size in tissue samples and may fail to capture cell size heterogeneity, an important pathological feature. Additionally, the correlation between the genes we screened and cell size does not equate to causality. Further studies combining molecular biology experiments are needed to clarify the regulatory relationship between them. As a basic morphological parameter, cell size also changes in many physiological or pathological processes, such as the menstrual cycle, pregnancy, and myocardial hypertrophy. The validity and application value of CSS under these conditions need exploration. In summary, the CSS developed in our study bridges gene expression and cell morphology. It not only reveals the critical role of cell size in processes such as cancer drug resistance, aging, and exercise adaptation, but also provides a new paradigm for future utilization of transcriptomic data to inspect cellular morphological features.

## Methods and Materials

### Construction of CSS

This study used publicly available IF images from the HPA database^20^ as the discovery dataset for obtaining cell sizes. The HPA IF images were acquired through a standardized immunocytochemistry (ICC) workflow^19^. Specifically, human cell lines were cultured on glass coverslips, fixed with paraformaldehyde, and permeabilized with Triton X-100^62^. Indirect IF staining was performed using primary antibodies against specific proteins and corresponding fluorescently labeled secondary antibodies, with multiplex labeling using DAPI (nucleus), anti-α-tubulin (microtubules), and anti-Calreticulin (endoplasmic reticulum). Images were acquired using a confocal laser scanning microscope equipped with a 63× oil immersion objective, ensuring high resolution and clear presentation of subcellular structures, which provided a high-quality data foundation for precise cell area measurement based on image segmentation. According to the HPA guidelines for programmatic access (https://www.proteinatlas.org/about/download), we extracted XML files containing information of the URLs and scale bar for the IF images. A total of 100,080 accessible images involving 42 cell lines were available. Additional filtering was performed: (1) images must contain scale bar information (μm/pixel); (2) the cell lines used must have expression profile data in the CCLE; (3) official cell segmentation mask images must be provided. As a result, 64,006 images remained.

For each IF image, we employed the connected component algorithm from OpenCV-Python (v4.13.0.92) to calculate cell areas. To avoid underestimation of area caused by cell truncation at image edges, we excluded all connected regions touching the image boundaries. Using the scale bar information from the images, pixel areas were converted to physical areas (μm²). Additionally, regions with areas smaller than 10 μm² were excluded to eliminate possible non-cellular structures (e.g., artifacts from cell segmentation, cell debris, etc.). Finally, a total of 519,632 cells from 63,475 IF images across 22 cell lines were used to evaluate cell size. For each cell line, we used the median of its areas as the metric for cell size.

The transcriptomic data for the above cells were obtained from the CCLE^21^ (https://sites.broadinstitute.org/ccle) expression matrix standardized by transcripts per million (TPM). For the 57,820 genes therein, we calculated the Spearman rank correlation coefficient (ρ) and p-value between the expression level of each gene and cell size. Subsequently, the Benjamini-Hochberg (BH) method was used to correct p-values for multiple testing under each metric, calculating the FDR. Finally, we selected the gene set positively correlated with cell area with FDR < 0.05 as the constituent genes of CSS, comprising a total of 457 genes (Supplementary Table S2).

For bulk transcriptomic data, CSS was calculated using the single sample Gene Set Enrichment Analysis (ssGSEA) algorithm. For single-cell and spatial transcriptomics, CSS was calculated using the AUCell algorithm. Both ssGSEA^63^ and AUCell^64^ are unsupervised scoring methods capable of evaluating the enrichment level of specific gene sets in individual samples/cells based on gene expression profiles. They are both rank-based enrichment analysis methods, with AUCell designed to better handle the sparsity characteristics of single-cell data. ssGSEA was implemented using the gseapy package^65^ (v1.1.13), with the normalized enrichment score (NES) serving as CSS. AUCell was obtained from decoupler^66^ (v2.1.6), with the AUC value serving as CSS.

### Acquisition and Processing of Validation Sets

We manually collected 37 reports on human cell line sizes from published literature and public databases^26–30^, involving 21 cell lines (Supplementary Table S3). Most data were recorded in the form of cell diameter (μm). For cells recorded in terms of volume, we approximated their diameters using the sphere volume formula. These data originated from different research backgrounds and experimental conditions, thus exhibiting high heterogeneity. We matched these cell lines with CCLE cell line IDs and performed correlation analysis between the corresponding CSS and the cell diameters reported in the literature to validate the accuracy of CSS.

For the mouse validation set, we obtained expression profiles of mouse hepatocytes with 4 different relative nuclear radii (positively correlated with cell size) from the supplementary materials of the study by Miettinen et al.^32^ (4 samples per group). Using BioMart^67^, we converted the 457 human cell size-related genes to mouse orthologs and calculated CSS using the same strategy (ssGSEA). Finally, correlation analysis was performed with the relative nuclear radius reported in the literature.

To validate CSS at single-cell resolution, we used cervical cancer spatial transcriptome data based on the Xenium platform officially provided by 10× Genomics (https://www.10xgenomics.com/datasets/xenium-prime-ffpe-human-cervical-cancer). Data preprocessing included: (1) excluding cells with a negative control probe ratio ≥ 2.0% to ensure robust signal-to-noise ratios; (2) excluding cells with total transcript counts below the 2.5th percentile or above the 97.5th percentile; (3) retaining only cells detecting no fewer than 10 different genes to exclude non-specific probe aggregation or cell debris; (4) excluding cells with areas below the 2.5th percentile or above the 97.5th percentile to avoid cell segmentation artifacts or doublets; (5) excluding cells segmented by nucleus expansion of 5.0µm, as their cell areas may be inaccurate. A total of 732,704 high-quality cells were finally obtained for validation.

### Correlation Analysis of CSS with Drug Sensitivity

Drug sensitivity data came from CTRP v2.0 (https://portals.broadinstitute.org/ctrp). CTRP provides dose-response data for large-scale cancer cell lines against hundreds of small molecule compounds^40^. We downloaded AUC values as the metric for drug sensitivity, where larger AUC indicates greater cell resistance to the drug. The CSS calculated from CCLE was merged with the AUC data from CTRP. For each compound, the Spearman correlation coefficient (ρ) between CSS and AUC was calculated, and p-value was corrected using the BH method.

Single-cell transcriptomic profiles paired with drug response phenotypes were obtained from scDrugAtlas^68^ (http://drug.hliulab.tech/scDrugAtlas). Three datasets were analyzed: dataset ID 95^69^ (Gemcitabine response), 66^70^ (Cisplatin response), and 83^71^ (Paclitaxel and Doxorubicin response). For UMAP visualization, the top 2,000 highly variable genes were identified and used for principal component analysis (PCA). The optimal number of principal components was determined by the Kneedle algorithm (kneed^72^ v0.8.6), which identifies the knee point (point of maximum curvature) in the cumulative variance curve.

### Survival Analysis

TCGA^11^ transcriptome (TPM normalized) and clinical data were obtained from the Genomic Data Commons (GDC) Data Portal (https://portal.gdc.cancer.gov). We calculated CSS for each sample, involving 33 cancer types. Only primary tumors (for solid tumors) and primary blood-derived cancers (for hematological cancers) were retained for analysis. After z-score standardization, CSS was included as a continuous variable in the Cox proportional hazards regression model (implemented from the Python package lifelines^73^ v0.30.3) to calculate the hazard ratio (HR) and its 95% confidence interval.

### Correlation Analysis of CSS with Age

RNA-seq data and sample/donor annotation data from version 8 of the GTEx Project^42^ were obtained from https://gtexportal.org/home/downloads/adult-gtex. We used the expression matrix (TPM normalized) to calculate CSS and performed the Spearman correlation analysis between CSS and donor age.

We integrated five scRNA-seq datasets of human ovary: GSE184880^74^, GSE255690^75^, GSE260685^44^, GSE285362^76^ from Gene Expression Omnibus (GEO, https://www.ncbi.nlm.nih.gov/gds), and ovarian data from the Tabula Sapiens project^12,77^ (v2, https://figshare.com/articles/dataset/Tabula_Sapiens_v2/27921984). Sample metadata, including donor age, were obtained from GEO or the original publications. For GSE184880, only normal tissue samples were retained; for GSE255690, spatial transcriptomics samples were excluded; for GSE285362, only fresh samples were retained. Quality control was performed per sample: Scrublet was used to detect and remove doublets; cells expressing fewer than 250 genes, with fewer than 400 total UMI counts, or with mitochondrial gene proportion exceeding 20% were filtered out. All samples were then concatenated and genes expressed in fewer than 3 cells were removed. As a result, 213,882 cells from 23 samples (ages 18∼56) were used for data integration. The top 2,000 highly variable genes were identified using the Seurat v3 flavor with batch key set to sample ID. A single-cell variational inference (scVI)^78^ variational autoencoder (scvi-tools v1.4.3, 2 hidden layers, 30 latent dimensions, negative binomial likelihood) was trained on these genes with sample ID as the batch key. The resulting latent representation was used for neighborhood graph construction, Leiden clustering, and UMAP dimensionality reduction (Supplementary Figure S3A). Cell type annotation (Supplementary Figure S3C) for each Leiden cluster comprehensively considered both consensus marker genes (Supplementary Figure S3B) and the neighborhood status in UMAP. For immune cells, we extracted T/NK cells, macrophages/monocytes, mast cells, and B/plasma cells, and performed annotation using CellTypist^79^ (v1.7.1) with the “Immune_All_High” model (Supplementary Figure S3D). Predictions with confidence above 99% were accepted. Cell types with fewer than 100 cells, as well as doublets (Leiden cluster 17), were excluded, resulting in 209,188 cells (**Figure 4J**). CSS was calculated per cell after library-size normalization (to 10,000) and log-transformation. To evaluate the association between CSS and age for each cell type, we fitted a linear regression model (statsmodels v0.14.6) with z-score standardized CSS as the dependent variable, donor age (scaled by dividing by 10) as the primary independent variable, and dataset as a categorical batch covariate (one-hot encoded). Robust standard errors (HC3 type) were used to account for potential heteroskedasticity.

### Physical Exercise Datasets

We collected 10 independent GEO datasets (GSE27536^80^, GSE28422^81^, GSE28998^82^, GSE47881^83^, GSE71972^84^, GSE72462^85^, GSE163356^86^, GSE182494^87^, GSE195585^88^, and GSE202295^89^), all uniformly organized from ExerGeneDB^90^ (https://exergenedb.com), containing transcriptomic data of skeletal muscle tissue before and after exercise intervention. Exercise intervention protocols included physical exercise of different intensities, durations, and types (endurance training and resistance training).

### GO Enrichment Analysis

We used the enrich function of the gseapy package^65^ (v1.1.13) for over-representation analysis (ORA). GO biological process gene sets^91^ were obtained from the Molecular Signatures Database^92^ (MSigDB, https://www.gsea-msigdb.org/gsea/msigdb, v2025.1). The target gene set was genes constituting CSS (457 positively correlated genes), and the background gene set was all detected genes in CCLE.

### Statistical Analysis

All statistical analyses were conducted within a Python (v3.13.12) environment. The study utilized SciPy^93^ (v1.17.1) for fundamental statistical testing. Two-sided Spearman rank correlation tests were used to assess the correlation between continuous variables or between continuous and ordinal categorical variables. Two-sided Wilcoxon rank-sum test was used for non-parametric comparison between two groups of continuous variables.

## Data Availability

All data generated or analyzed during this study are included in this article and its supplementary information files. The discovery dataset consisted of IF images obtained from the HPA database (https://www.proteinatlas.org/about/download), with corresponding transcriptomic profiles from the CCLE (https://sites.broadinstitute.org/ccle). Other datasets were sourced from multiple public repositories: cell size measurements from literature reports (Supplementary Table S3), mouse liver cell data from Miettinen et al.^32^, Xenium spatial transcriptomics data from 10x Genomics (https://www.10xgenomics.com/datasets/xenium-prime-ffpe-human-cervical-cancer), drug sensitivity data from the CTRP (https://portals.broadinstitute.org/ctrp), scRNA-seq data (dataset ID: 66, 83, and 95) with drug response phenotypes from scDrugAtlas (http://drug.hliulab.tech/scDrugAtlas), transcriptomic and clinical survival data from TCGA via the GDC data portal (https://portal.gdc.cancer.gov), aging-related transcriptomic data from the GTEx Project (https://gtexportal.org/home/downloads/adult-gtex), ovarian scRNA-seq data (GSE184880, GSE255690, GSE260685, and GSE285362) from GEO (https://www.ncbi.nlm.nih.gov/gds), ovarian scRNA-seq data from Tabula Sapiens project (https://figshare.com/articles/dataset/Tabula_Sapiens_v2/27921984), and exercise intervention transcriptomic datasets (GSE27536, GSE28422, GSE28998, GSE47881, GSE71972, GSE72462, GSE163356, GSE182494, GSE195585, and GSE202295) from ExerGeneDB (https://exergenedb.com). GO gene sets were obtained from MSigDB (https://www.gsea-msigdb.org/gsea/msigdb). The complete list of 457 genes constituting the CSS is provided in Supplementary Table S2. All data generated in this study are available from the corresponding author upon reasonable request.

## Code Availability

The code used for calculating CSS is available at https://github.com/Evan210/cell_size_score.

## Acknowledgments

This study was partly supported by the grants from the National Natural Science Foundation of China (No. 62501021, 82427801) and the China Postdoctoral Science Foundation (No. BX20240027). We would like to thank the creators on SciDraw (https://scidraw.io) for providing the scientific illustrations used in Figure 1A, specifically John Chilton (doi: 10.5281/zenodo.3926101, 10.5281/zenodo.3926109, 10.5281/zenodo.3926133, 10.5281/zenodo.3926161, 10.5281/zenodo.3926211, 10.5281/zenodo.3926221, 10.5281/zenodo.3926261, 10.5281/zenodo.3926367, 10.5281/zenodo.3926537, 10.5281/zenodo.3926549, and 10.0.20.161/zenodo.14284999) and an anonymous artist (doi: 10.0.20.161/zenodo.14059328). We appreciate the authors of the studies for making their data publicly available and all the donors who contributed to the research.

## Author contributions

Q.C. conceived the project and thoroughly revised the manuscript. X.J. performed the study and wrote the raw manuscript. Q.C. supervised the study. All authors read and approved the final manuscript.

## References

1. Greenough, R. B. Varying Degrees of Malignancy in Cancer of the Breast. J. Cancer Res. 9, 453–463 (1925).

2. Herbein, G. & Nehme, Z. Polyploid Giant Cancer Cells, a Hallmark of Oncoviruses and a New Therapeutic Challenge. Front. Oncol. 10, 567116 (2020).

3. Lanz, M. C. et al. Increasing cell size remodels the proteome and promotes senescence. Mol. Cell 82, 3255–3269.e8 (2022).

4. Itgen, M. W., Natalie, G. R., Siegel, D. S., Sessions, S. K. & Mueller, R. L. Genome size drives morphological evolution in organ-specific ways. Evol. Int. J. Org. Evol. 76, 1453–1468 (2022).

5. Scepanovic, G. & Fernandez-Gonzalez, R. Should I shrink or should I grow: cell size changes in tissue morphogenesis. Genome 67, 125–138 (2024).

6. Seal, S. et al. Cell Painting: a decade of discovery and innovation in cellular imaging. Nat. Methods 22, 254–268 (2025).

7. Ramezani, M. et al. A genome-wide atlas of human cell morphology. Nat. Methods 22, 621–633 (2025).

8. Marco Salas, S., et al. Optimizing Xenium In Situ data utility by quality assessment and best-practice analysis workflows. Nat. Methods 22, 813–823 (2025).

9. Oliveira, M. F. de et al. High-definition spatial transcriptomic profiling of immune cell populations in colorectal cancer. Nat. Genet. 57, 1512–1523 (2025).

10. Wei, X. et al. Single-cell Stereo-seq reveals induced progenitor cells involved in axolotl brain regeneration. Science 377, eabp9444 (2022).

11. Cancer Genome Atlas Research Network et al. The Cancer Genome Atlas Pan-Cancer analysis project. Nat. Genet. 45, 1113–1120 (2013).

12. Tabula Sapiens Consortium* et al. The Tabula Sapiens: A multiple-organ, single-cell transcriptomic atlas of humans. Science 376, eabl4896 (2022).

13. Fingar, D. C., Salama, S., Tsou, C., Harlow, E. & Blenis, J. Mammalian cell size is controlled by mTOR and its downstream targets S6K1 and 4EBP1/eIF4E. Genes Dev. 16, 1472–1487 (2002).

14. Mafi, S. et al. mTOR-Mediated Regulation of Immune Responses in Cancer and Tumor Microenvironment. Front. Immunol. 12, 774103 (2021).

15. Conciatori, F. et al. mTOR Cross-Talk in Cancer and Potential for Combination Therapy. Cancers 10, 23 (2018).

16. Padovan-Merhar, O. et al. Single mammalian cells compensate for differences in cellular volume and DNA copy number through independent global transcriptional mechanisms. Mol. Cell 58, 339–352 (2015).

17. Chen, Y., Zhao, G., Zahumensky, J., Honey, S. & Futcher, B. Differential Scaling of Gene Expression with Cell Size May Explain Size Control in Budding Yeast. Mol. Cell 78, 359–370.e6 (2020).

18. Jones, I. et al. Characterization of proteome-size scaling by integrative omics reveals mechanisms of proliferation control in cancer. Sci. Adv. 9, eadd0636 (2023).

19. Barbe, L. et al. Toward a confocal subcellular atlas of the human proteome. Mol. Cell. Proteomics MCP 7, 499–508 (2008).

20. Thul, P. J. et al. A subcellular map of the human proteome. Science 356, eaal3321 (2017).

21. Ghandi, M. et al. Next-generation characterization of the Cancer Cell Line Encyclopedia. Nature 569, 503–508 (2019).

22. Zhang, J., Liu, Y., Guo, Y. & Zhao, Q. GPX8 promotes migration and invasion by regulating epithelial characteristics in non-small cell lung cancer. Thorac. Cancer 11, 3299–3308 (2020).

23. Wang, L. et al. Regulators of tubulin polyglutamylation control nuclear shape and cilium disassembly by balancing microtubule and actin assembly. Cell Res. 32, 190–209 (2022).

24. Qin, L. et al. SNX7 mediates inhibition of autophagy in prostate cancer via activation of CFLIP expression. Discov. Oncol. 16, 1656 (2025).

25. Xu, L. et al. An antiapoptotic role of sorting nexin 7 is required for liver development in zebrafish. Hepatology 55, 1985–1993 (2012).

26. Khetan, J., Shahinuzzaman, M., Barua, S. & Barua, D. Quantitative Analysis of the Correlation between Cell Size and Cellular Uptake of Particles. Biophys. J. 116, 347–359 (2019).

27. Wang, M. et al. The effect of substrate stiffness on cancer cell volume homeostasis. J. Cell. Physiol. 233, 1414–1423 (2018).

28. Zhou, B., Lu, X., Hao, Y. & Yang, P. Real-Time Monitoring of the Regulatory Volume Decrease of Cancer Cells: A Model for the Evaluation of Cell Migration. Anal. Chem. 91, 8078–8084 (2019).

29. Achilles, K. & Bednarski, P. J. Quantification of elastase-like activity in 13 human cancer cell lines and in an immortalized human epithelial cell line by RP-HPLC. Biol. Chem. 384, 817–824 (2003).

30. Milo, R., Jorgensen, P., Moran, U., Weber, G. & Springer, M. BioNumbers--the database of key numbers in molecular and cell biology. Nucleic Acids Res. 38, D750–753 (2010).

31. Wagner, B. A., Venkataraman, S. & Buettner, G. R. The rate of oxygen utilization by cells. Free Radic. Biol. Med. 51, 700–712 (2011).

32. Miettinen, T. P. et al. Identification of transcriptional and metabolic programs related to mammalian cell size. Curr. Biol. CB 24, 598–608 (2014).

33. Yu, F.-X., Zhao, B. & Guan, K.-L. Hippo Pathway in Organ Size Control, Tissue Homeostasis, and Cancer. Cell 163, 811–828 (2015).

34. Dupont, S. et al. Role of YAP/TAZ in mechanotransduction. Nature 474, 179–183 (2011).

35. Hoxhaj, G. & Manning, B. D. The PI3K-AKT network at the interface of oncogenic signalling and cancer metabolism. Nat. Rev. Cancer 20, 74–88 (2020).

36. Saxton, R. A. & Sabatini, D. M. mTOR Signaling in Growth, Metabolism, and Disease. Cell 168, 960–976 (2017).

37. Zhou, M., Zhou, M. & Jin, Y. Tumour Cell Size Control and Its Impact on Tumour Cell Function. Cell Prolif. 58, e70080 (2025).

38. Pettersen, A. K., Schuster, L. & Metcalfe, N. B. The Evolution of Offspring Size: a Metabolic Scaling Perspective. Integr. Comp. Biol. 62, 1492–1502 (2022).

39. Scriven, J. J., Whitehorn, P. R., Goulson, D. & Tinsley, M. C. Bergmann’s Body Size Rule Operates in Facultatively Endothermic Insects: Evidence from a Complex of Cryptic Bumblebee Species. PloS One 11, e0163307 (2016).

40. Seashore-Ludlow, B. et al. Harnessing Connectivity in a Large-Scale Small-Molecule Sensitivity Dataset. Cancer Discov. 5, 1210–1223 (2015).

41. Becht, E. et al. Dimensionality reduction for visualizing single-cell data using UMAP. Nat. Biotechnol. (2018) doi:10.1038/nbt.4314.

42. GTEx Consortium. The GTEx Consortium atlas of genetic regulatory effects across human tissues. Science 369, 1318–1330 (2020).

43. Blagosklonny, M. V. Cell senescence, rapamycin and hyperfunction theory of aging. Cell Cycle 21, 1456–1467 (2022).

44. Jones, A. S. K. et al. Cellular atlas of the human ovary using morphologically guided spatial transcriptomics and single-cell sequencing. Sci. Adv. 10, eadm7506 (2024).

45. Zhang, Z., Huang, L. & Brayboy, L. Macrophages: an indispensable piece of ovarian health. Biol. Reprod. 104, 527–538 (2021).

46. Zhang, Z., Schlamp, F., Huang, L., Clark, H. & Brayboy, L. Inflammaging is associated with shifted macrophage ontogeny and polarization in the aging mouse ovary. Reproduction 159, 325–337 (2020).

47. Schorr, M. et al. Sex differences in body composition and association with cardiometabolic risk. Biol. Sex Differ. 9, 28 (2018).

48. Martin, T. G. & Leinwand, L. A. Hearts apart: sex differences in cardiac remodeling in health and disease. J. Clin. Invest. 134, e180074 (2024).

49. Triebner, K. et al. Menopause Is Associated with Accelerated Lung Function Decline. Am. J. Respir. Crit. Care Med. 195, 1058–1065 (2017).

50. Haq, I. I., Banerjee, A. A., Arshad, Z., Iqbal, A. M. & Bhatia, M. The management of tarsal tunnel syndrome: A scoping review. J. Clin. Orthop. Trauma 54, 102489 (2024).

51. Ojkic, N., Serbanescu, D. & Banerjee, S. Antibiotic Resistance via Bacterial Cell Shape-Shifting. mBio 13, e0065922 (2022).

52. Neurohr, G. E. et al. Excessive Cell Growth Causes Cytoplasm Dilution And Contributes to Senescence. Cell 176, 1083–1097.e18 (2019).

53. Coward, J. & Harding, A. Size Does Matter: Why Polyploid Tumor Cells are Critical Drug Targets in the War on Cancer. Front. Oncol. 4, 123 (2014).

54. Huang, P. et al. Polyploid giant cancer cells and tumor budding: translation from basic research to clinical application. Front. Oncol. 15, 1611920 (2025).

55. Zhang, X. et al. Targeting polyploid giant cancer cells potentiates a therapeutic response and overcomes resistance to PARP inhibitors in ovarian cancer. Sci. Adv. 9, eadf7195 (2023).

56. Ma, Y. et al. High-Throughput Empirical and Virtual Screening To Discover Novel Inhibitors of Polyploid Giant Cancer Cells in Breast Cancer. Anal. Chem. 97, 5498–5506 (2025).

57. Pianka, E. R. On r-and K-selection. Am. Nat. 104, 592–597 (1970).

58. Sender, R. & Milo, R. The distribution of cellular turnover in the human body. Nat. Med. 27, 45–48 (2021).

59. Hatton, I. A. et al. The human cell count and size distribution. Proc. Natl. Acad. Sci. U. S. A. 120, e2303077120 (2023).

60. Schoenfeld, B. J. Does exercise-induced muscle damage play a role in skeletal muscle hypertrophy? J. Strength Cond. Res. 26, 1441–1453 (2012).

61. Schoenfeld, B. J. & Contreras, B. The muscle pump: potential mechanisms and applications for enhancing hypertrophic adaptations. Strength Cond. J. 36, 21–25 (2014).

62. Stadler, C., Skogs, M., Brismar, H., Uhlén, M. & Lundberg, E. A single fixation protocol for proteome-wide immunofluorescence localization studies. J. Proteomics 73, 1067–1078 (2010).

63. Barbie, D. A. et al. Systematic RNA interference reveals that oncogenic KRAS-driven cancers require TBK1. Nature 462, 108–112 (2009).

64. Aibar, S. et al. SCENIC: single-cell regulatory network inference and clustering. Nat. Methods 14, 1083–1086 (2017).

65. Fang, Z., Liu, X. & Peltz, G. GSEApy: a comprehensive package for performing gene set enrichment analysis in Python. Bioinformatics 39, btac757 (2023).

66. Badia-I-Mompel, P., et al. decoupleR: ensemble of computational methods to infer biological activities from omics data. Bioinforma. Adv. 2, vbac016 (2022).

67. Smedley, D. et al. BioMart--biological queries made easy. BMC Genomics 10, 22 (2009).

68. Wu, Y., Huang, W., Ren, X., Liu, H. & Xu, L. scDrugAtlas: an integrative single-cell drug response database for dissecting tumour heterogeneity in therapeutic efficacy. Database J. Biol. Databases Curation 2026, baag010 (2026).

69. Principe, D. R. et al. Calcium channel blockers potentiate gemcitabine chemotherapy in pancreatic cancer. Proc. Natl. Acad. Sci. U. S. A. 119, e2200143119 (2022).

70. Stewart, C. A. et al. Single-cell analyses reveal increased intratumoral heterogeneity after the onset of therapy resistance in small-cell lung cancer. Nat. Cancer 1, 423–436 (2020).

71. Stevens, L. E. et al. JAK-STAT Signaling in Inflammatory Breast Cancer Enables Chemotherapy-Resistant Cell States. Cancer Res. 83, 264–284 (2023).

72. Satopaa, V., Albrecht, J., Irwin, D. & Raghavan, B. Finding a ‘Kneedle’ in a Haystack: Detecting Knee Points in System Behavior. in 2011 31st International Conference on Distributed Computing Systems Workshops 166–171 (2011). doi:10.1109/ICDCSW.2011.20.

73. Davidson-Pilon, C. lifelines: survival analysis in Python. J. Open Source Softw. 4, 1317 (2019).

74. Xu, J. et al. Single-Cell RNA Sequencing Reveals the Tissue Architecture in Human High-Grade Serous Ovarian Cancer. Clin. Cancer Res. Off. J. Am. Assoc. Cancer Res. 28, 3590–3602 (2022).

75. Wu, M. et al. Spatiotemporal transcriptomic changes of human ovarian aging and the regulatory role of FOXP1. *Nat*. Aging 4, 527–545 (2024).

76. Guo, F. et al. Identification of cryosensitive niches and a targetable FOS/AP-1 program in the human ovarian cortex by single-cell and spatial transcriptomics. BMC Med. 24, 206 (2026).

77. Quake, S. R. & The Tabula Sapiens Consortium. Tabula Sapiens reveals transcription factor expression, senescence effects, and sex-specific features in cell types from 28 human organs and tissues. bioRxiv 2024.12.03.626516 (2024) doi:10.1101/2024.12.03.626516.

78. Lopez, R., Regier, J., Cole, M. B., Jordan, M. I. & Yosef, N. Deep generative modeling for single-cell transcriptomics. Nat. Methods 15, 1053–1058 (2018).

79. Domínguez Conde, C., et al. Cross-tissue immune cell analysis reveals tissue-specific features in humans. Science 376, eabl5197 (2022).

80. Turan, N. et al. A systems biology approach identifies molecular networks defining skeletal muscle abnormalities in chronic obstructive pulmonary disease. PLoS Comput. Biol. 7, e1002129 (2011).

81. Raue, U. et al. Transcriptome signature of resistance exercise adaptations: mixed muscle and fiber type specific profiles in young and old adults. J. Appl. Physiol. 112, 1625–1636 (2012).

82. Gordon, P. M. et al. Resistance exercise training influences skeletal muscle immune activation: a microarray analysis. J. Appl. Physiol. 112, 443–453 (2012).

83. Phillips, B. E. et al. Molecular networks of human muscle adaptation to exercise and age. PLoS Genet. 9, e1003389 (2013).

84. Romero, S. A. et al. Evidence of a broad histamine footprint on the human exercise transcriptome. J. Physiol. 594, 5009–5023 (2016).

85. Böhm, A. et al. TGF-β Contributes to Impaired Exercise Response by Suppression of Mitochondrial Key Regulators in Skeletal Muscle. Diabetes 65, 2849–2861 (2016).

86. Norrbom, J. M. et al. A HIF-1 signature dominates the attenuation in the human skeletal muscle transcriptional response to high-intensity interval training. J. Appl. Physiol. 132, 1448–1459 (2022).

87. Dial, A. G. et al. Alterations in skeletal muscle repair in young adults with type 1 diabetes mellitus. Am. J. Physiol. Cell Physiol. 321, C876–C883 (2021).

88. Centner, C. et al. Supplementation of Specific Collagen Peptides Following High-Load Resistance Exercise Upregulates Gene Expression in Pathways Involved in Skeletal Muscle Signal Transduction. Front. Physiol. 13, 838004 (2022).

89. Pillon, N. J. et al. Distinctive exercise-induced inflammatory response and exerkine induction in skeletal muscle of people with type 2 diabetes. Sci. Adv. 8, eabo3192 (2022).

90. Pan, L. et al. ExerGeneDB: A physical exercise-regulated differential gene expression database. J. Sport Health Sci. 14, 101027 (2025).

91. Gene Ontology Consortium et al. The Gene Ontology knowledgebase in 2023. Genetics 224, iyad031 (2023).

92. Liberzon, A. et al. The Molecular Signatures Database (MSigDB) hallmark gene set collection. Cell Syst. 1, 417–425 (2015).

93. Virtanen, P. et al. SciPy 1.0: fundamental algorithms for scientific computing in Python. Nat. Methods 17, 261–272 (2020).

