## Supplementary Figures for "Bridging Gene Expression and Morphology: A Cell Size Score and Its Applications Across Multiple Diseases and Physiological Contexts"

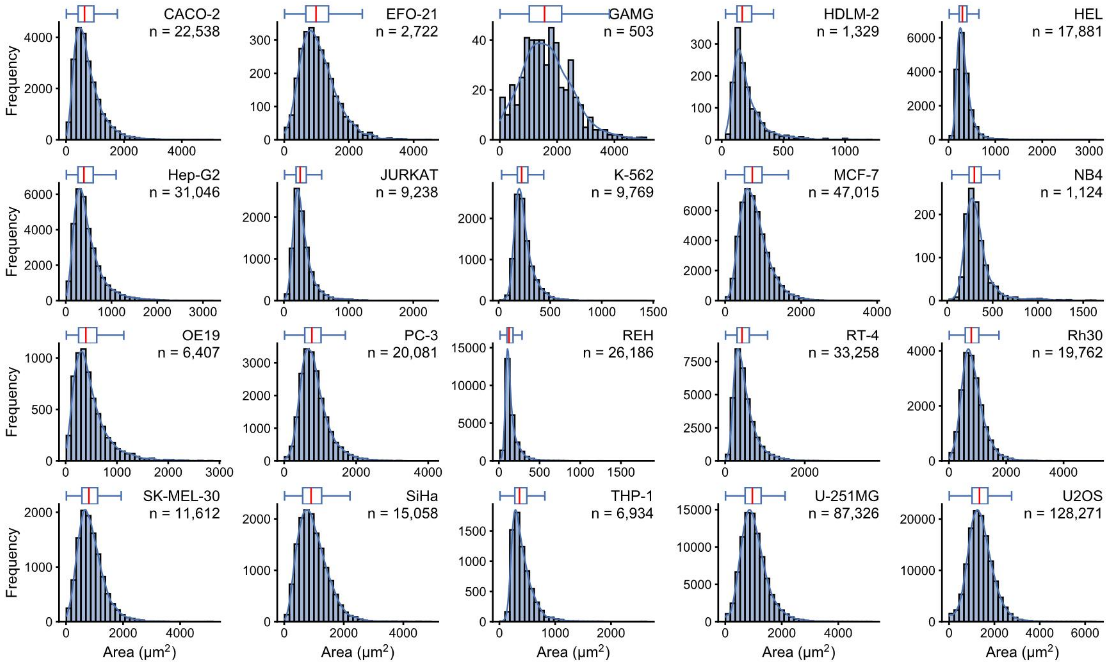


**Supplementary Figure S1. Cell area distribution profiles across 20 distinct cell lines.** Histograms and kernel density estimation (KDE) curves illustrating the distribution of cell area (μm²) for 20 different human cell lines. The horizontal box plots above each histogram summarize the descriptive statistics of the population: the center line denotes the median area, the left and right edges of the blue box indicate the 25th and 75th percentiles, respectively, and the whiskers extend to the most extreme values within 1.5 times the interquartile range.

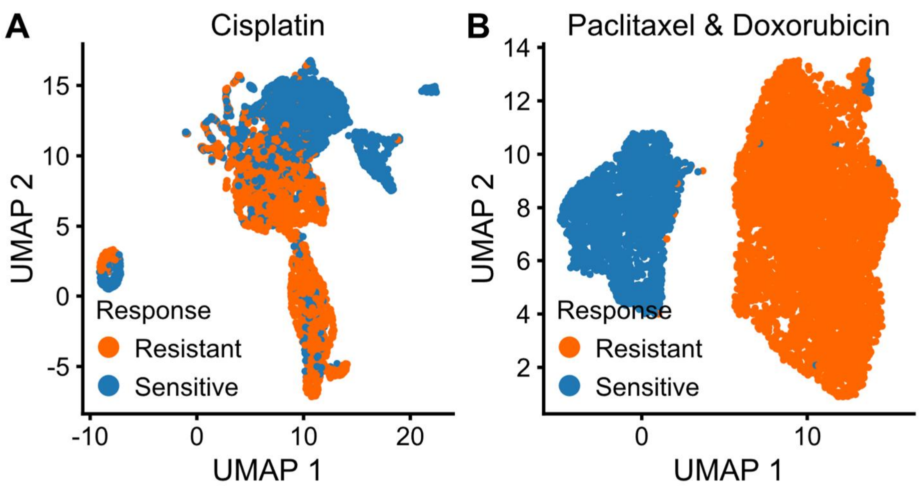


**Supplementary Figure S2. Single cell UMAP visualization of cell clustering based on drug response.**

(A-B) UMAP visualization of cell clustering based on (A) Cisplatin and (B) Paclitaxel & Doxorubicin drug response.

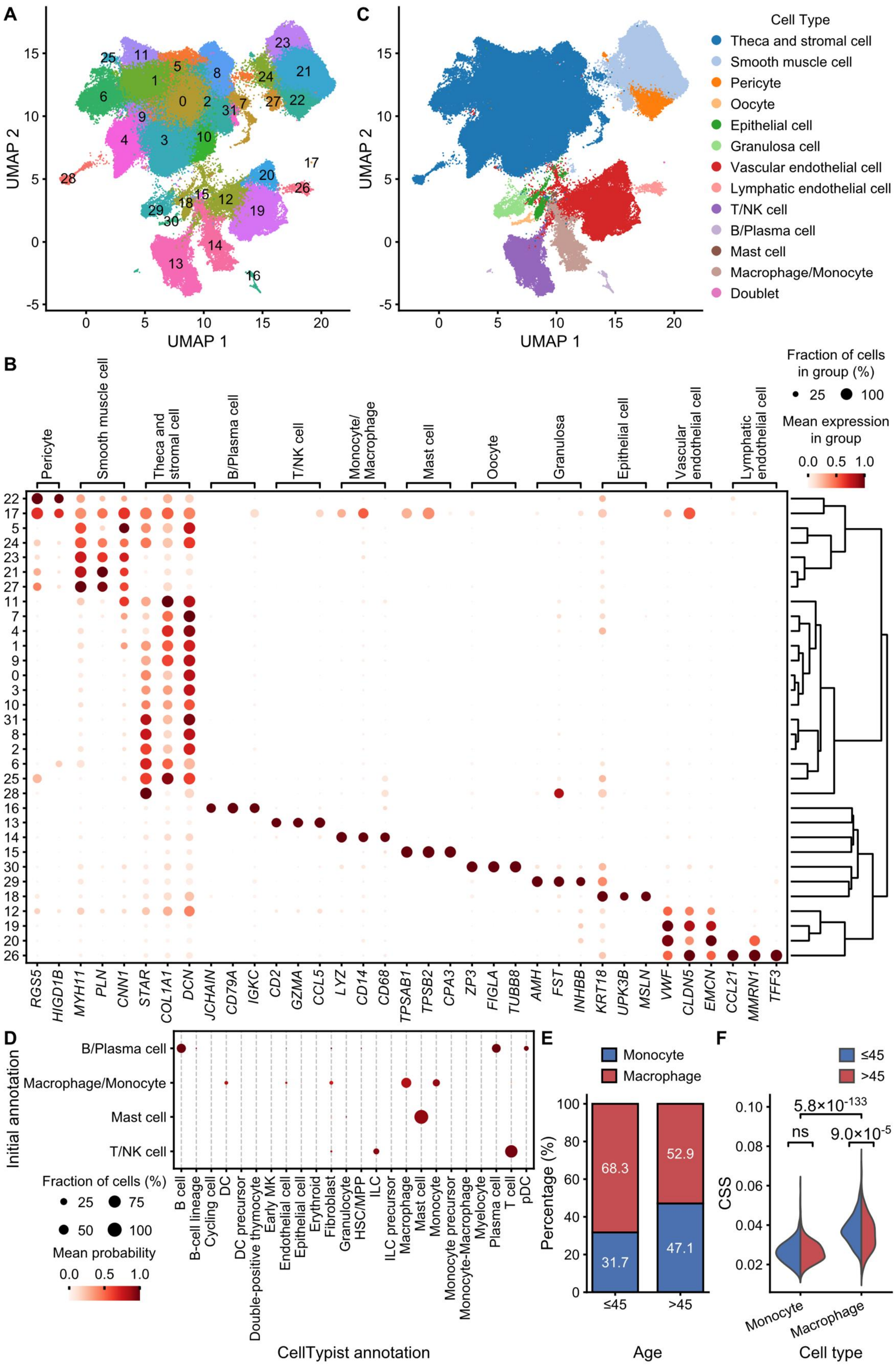


**Supplementary Figure S3. Single-cell analysis of aging ovaries.**

(Legend continued on next page)

**Supplementary Figure S3. Single-cell analysis of aging ovaries.**

(A) UMAP visualization of all ovarian cells colored by Leiden clusters.

(B) Dot plot showing the expression of marker genes for major cell types. Dot size represents the percentage of cells expressing the gene, and color intensity represents the mean expression level.

(C) UMAP visualization of ovarian cells colored by initially annotated cell types.

(D) Dot plot showing the cell type annotation by CellTypist. Dot size represents the proportion of initially annotated cell types predicted by CellTypist as various cell types, while the color intensity indicates the average predicted probability. DC: dendritic cell; Early MK: early megakaryocyte; HSC/MPP: hematopoietic stem cell/multipotent progenitor; ILC: innate lymphoid cell; pDC: plasmacytoid dendritic cell.

(E) Stacked bar chart showing the proportion of monocytes and macrophages in different age groups (≤45 vs. >45).

(F) Violin plot comparing the Cell Size Score (CSS) between monocytes and macrophages, split by age groups (≤45 vs. >45). Two-sided Wilcoxon's rank sum tests are used, ns: not significant ($p \geq 0.05$).

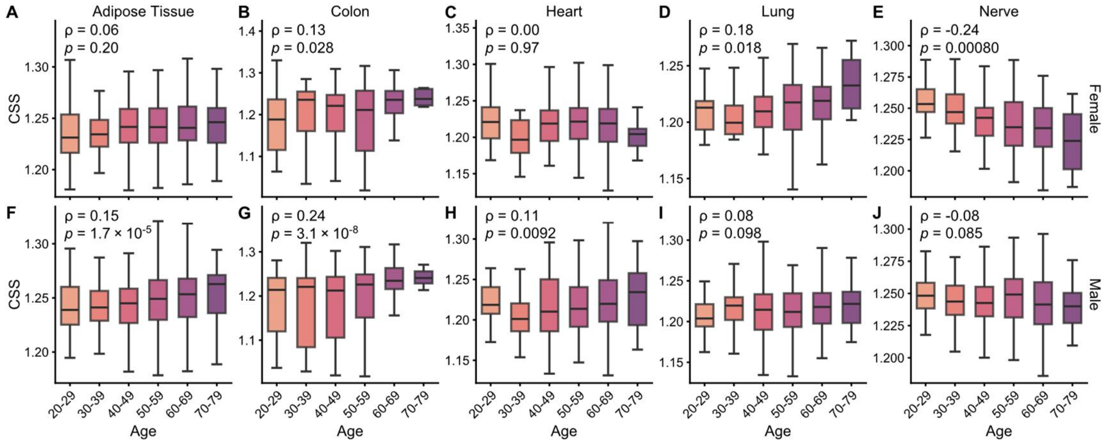

**Supplementary Figure S4. Sex stratification of age-dependent changes of cell size.**

The distribution of Cell Size Score (CSS) across age decades (20-29 to 70-79 years) between (A-E, top row) female and (F-J, bottom row) male. Tissues analyzed include (A, F) adipose tissue, (B, G) colon, (C, H) heart, (D, I) lung, and (E, J) nerve. The center line represents the median, box limits represent the 25th and 75th percentiles, respectively, and the whiskers extend to the most extreme values within 1.5 times the interquartile range. Spearman correlation coefficients (ρ) and p values are annotated.
